# High Consumption of Coffee Disrupts Nonhomologous End Joining Implications for Genomic Stability

**DOI:** 10.64898/2026.04.05.716533

**Authors:** Susmita Kumari, Humaira Siddiqua, Sathees C. Raghavan

## Abstract

Caffeine, the most widely consumed stimulant worldwide and primarily sourced from coffee, is well known for its central nervous system effects. Emerging evidence indicates that caffeine also modulates key cellular processes, including DNA repair. It inhibits the kinase activity of ATM and ATR-essential DNA damage response proteins, and impairs homologous recombination (HR)-mediated repair through multiple mechanisms. However, its effects on nonhomologous end joining (NHEJ), a major double-strand break (DSB) repair pathway, have been underexplored. In a recent study, we reported that caffeine inhibits NHEJ primarily by interfering with Ligase IV/XRCC4 complex, using *in vitro* and *ex vivo* model systems. Given coffee’s role as a primary dietary caffeine source, this study investigates the impact of *Coffea arabica* decoction on NHEJ-mediated DSB repair. High-performance liquid chromatography (HPLC) quantified caffeine levels in the decoction, followed by *in vitro* and *ex vivo* assays to evaluate NHEJ efficiency. Results demonstrate that coffee decoction inhibits end joining of both compatible and noncompatible DNA ends in cell-free systems derived from normal and cancer cells. Extrachromosomal repair assays confirmed impaired intracellular NHEJ, leading to accumulation of unrepaired DSBs in human cells. Kinetic analysis of γ-H2AX foci formation and resolution revealed persistent DNA breaks and reduced repair kinetics. Reconstitution experiments verified that the decoction specifically targets the Ligase IV/XRCC4 complex. These findings, building on our previous work, establish coffee decoction as a potent NHEJ inhibitor, mirroring purified caffeine’s effects. This underscores caffeine’s interference with endogenous DNA repair, with profound implications for cancer therapy by sensitizing tumors to genotoxic treatments.

## Introduction

Caffeine is one of the world’s most commonly consumed dietary ingredients, found in coffee, tea, soft drinks, chocolate products, prescription drugs, and dietary supplements ^1, 2^. It occurs naturally in varying amounts in the leaves, fruits, and beans of over 60 plant species. Primary dietary sources include roasted coffee beans (*Coffea arabica* and *Coffea robusta*) and tea leaves (*Camellia sinensis*) ^3^. The other common sources of caffeine include the cacao bean (*Theobroma cacao*), kola nuts (*Cola acuminata*), Yaupon (*Ilex vomitoria*), guarana berries (*Paullinia cupana*), Guayusa (*Ilex guayusa*), and yerba mate (*Ilex paraguariensis*) ^2, 4^. Coffee beans also contain minor amounts of other methylxanthines, though at less than 1% of total methylxanthines ^5^.

About 80% of the global population consumes caffeinated products for their stimulating effects ^6, 7^. Over the past 50 years, global coffee consumption has grown at an average of 1.9% annually, reaching 9.7 million tons in 2018, with the highest levels in the Americas, Europe, and Japan ^8^. Intake varies widely by product type, influenced by factors like sex, age, geography, and cultural habits ^9^.

Beyond stimulation, caffeine, a methylxanthine and adenosine analog, inhibits ATM and ATR kinases, making it a key tool for studying DNA repair and cell cycle responses ^10, 11^. Double-strand breaks (DSBs) rank among the most severe DNA lesions, threatening genomic integrity and cell survival if unrepaired ^12, 13^. Mammalian cells employ multiple DSB repair pathways, including homologous recombination (HR), nonhomologous end joining (NHEJ), and microhomology-mediated end joining (MMEJ) ^14, 15, 16^. While its suppression of homologous recombination (HR) is well documented ^10, 11, 17, 18, 19^, effects on nonhomologous end joining (NHEJ), a major DSB repair pathway, were underexplored. High concentrations inhibit DNA-PKcs *in vitro* (Block et al., 2004; Wang et al., 2003), but *in vivo* impacts were unclear until our recent work showing caffeine impairs NHEJ and elevates chromosomal DSBs*. In silico*, biophysical, and biochemical analyses revealed that caffeine directly binds XRCC4, disrupting the crucial Ligase IV-mediated ligation step associated with the NHEJ ^20^. Given the coffee as the dominant dietary source, we examined *Coffea arabica* powder-derived coffee decoction for its effect on NHEJ. We quantified caffeine in *Coffea arabica* decoction via high-performance liquid chromatography (HPLC) for direct comparison with purified caffeine. We then assessed its effects on NHEJ efficiency using biochemical assays and *ex vivo* cellular reporter systems. Integrating mechanistic reconstitution and DNA damage/repair kinetics revealed a functional link between this dietary staple and NHEJ regulation. Habitual exposure may thus modulate DSB repair, with concentration-dependent outcomes; genomic instability in normal cells or sensitization of cancer cells to DNA-damaging therapies. These insights position coffee-derived caffeine as a candidate for enhancing cancer treatment via targeted NHEJ inhibition.

## Materials and Methods

### Enzymes, chemicals, and reagents

All chemicals and reagents were from Merck (USA), SRL (India), or HiMedia (India). DNA-modifying enzymes were from New England Biolabs (USA). Cell culture media were from Lonza (Switzerland) or MP Biomedicals (USA); foetal bovine serum and Pen-Strep from Gibco (USA). Radioisotope-labeled nucleotides were from Revvity (USA); antibodies from BD Bioscience (USA) or BioLegend (USA). Purified caffeine was from Merck (USA). *Coffea arabica* beans were sourced from Kerala farms (India), roasted, and ground into pure powder without additives.

### Oligomers

Oligomers used in the study is presented as Table S1.

### Mammalian cell culture

Human cell lines, Molt4 (acute lymphoblastic leukemia) and Jurkat (T cell leukemia) were purchased from National Center for Cell Science, Pune, India. HMF-3S (human mammary fibroblasts) were kindly gifted by Dr. Ramray Bhat (IISc, Bangalore). Cells were grown in DMEM or RPMI medium supplemented with 10% FBS and 100 μg/ml Penicillin G and Streptomycin. Cells were incubated at 37°C in a humidified atmosphere containing 5% CO_2_.

### Plasmids

The *in vivo* NHEJ reporter pimEJ5-GFP was a gift from Dr. Jeremy Stark (USA) ^21^. The I-SceI overexpression vector was from Dr. Ralph Scully (USA) ^22^. The Ligase IV/XRCC4 co-expression plasmid was a gift from Dr. Mauro Modesti (France) ^23^.

### Radiolabelled oligomeric DNA substrate preparation

5’-end labelling of oligomeric DNA was performed as described ^24, 25^ using T4 polynucleotide kinase in a buffer containing 20 mM Tris-acetate (pH 7.9), 10 mM magnesium acetate, 50 mM potassium acetate, 1 mM DTT, and [γ-³²P]ATP at 37°C for 1 h. Labelled products were purified through Sephadex G-25 columns (Sigma, USA) and stored at −20°C. Double-stranded oligomeric DNA substrates were prepared by annealing γ-³²P-labeled 75 nt oligomer SCR19 with unlabeled complementary 75 nt oligomers (10 mM NaCl, 1 mM EDTA) via boiling for 10 min followed by slow cooling: compatible 5’ overhangs (SCR19 + SCR20; non-compatible 5’-5’ overhangs (SCR19 + VK11); and 5’-3’ overhangs (SCR19 + VK13) ^26, 27^.

### HPLC analysis of caffeine in coffee decoction

Caffeine content in coffee decoctions was quantified by high-performance liquid chromatography (HPLC; Shimadzu, Kyoto, Japan) on a C18 reverse-phase column with UV/Vis detection at 272 nm (Fajara and Susanti, 2017; Karau et al., 2010). Standard caffeine (Sigma) solutions were prepared for calibration curve generation (peak area vs. concentration). The mobile phase was methanol:water (1:1). Decoction samples were analyzed by comparing retention times and spectral data to standards; concentrations were determined from peak areas using LabSolutions software (Shimadzu, Kyoto, Japan).

### Preparation of cell-free extract from mammalian cells

Cell-free extract (CFE) was prepared from Molt4, Jurkat and HMF-3S cells as described previously ^28, 29, 30, 31^. Approximately 3×10^7^ cells were resuspended in 2 volumes of hypotonic buffer (10 mM Tris-HCl [pH 8.0], 1 mM EDTA, and 5 mM DTT and protease inhibitors) followed by homogenization and incubation at 4°C for 20 min. Following this, half volume of high-salt buffer (50 mM Tris-HCl [pH 7.5], 1 M KCl, 2 mM EDTA and 2 mM DTT) was added, homogenized and incubated for 20 min at 4°C. Extracts were ultracentrifuged at 4°C for 3 h at 42,000 RPM in a TLA-100 rotor. Supernatant was collected and dialyzed in dialysis buffer (20 mM Tris-HCl [pH 8.0], 0.1 M potassium acetate, 20% v/v glycerol, 0.5 mM EDTA, 1 mM DTT and 0.1 mM PMSF) overnight. Extracts were aliquoted, snap frozen and stored at -80°C until further use.

### DNA end joining reactions using cell-free extracts

Extracts prepared from mammalian cancerous cell lines (Molt4, Jurkat and HMF-3S) (1 µg) were incubated with radiolabelled double-stranded (ds) DNA oligomers at 25°C in NHEJ buffer [30 mM HEPES-KOH (pH 7.9), 7.5 mM MgCl_2_, 1 mM DTT, 2 mM ATP, 50 μM dNTPs and 0.1 μg BSA] in the presence of increasing concentration of coffee decoction (1.25, 2.5, 5, 10, 20 and 40 mg/ml) ^32, 33^. The reaction mixture where the extract was not added served as the no protein control (NPC). The joining reaction was terminated by adding EDTA (10 mM). The DNA products were purified by phenol:chloroform extraction and precipitated with chilled ethanol and glycogen. The dried pellet was dissolved in 10 μl of 1X TE and resolved on 8% denaturing PAGE, depending on the size of the DNA substrate used. The gel was dried at 80°C for 45 min, exposed, and the signal was detected using phosphorImager FLA9000 (Fuji, Japan). MultiGauge software was used to quantify joined products in Photo Stimulated Luminescence Units (PSLU). The extent of inhibition in the caffeine-treated samples was similarly quantified in PSLU.

### Isolation of plasmid DNA

Plasmid DNA was isolated from *E. coli* DH5α by alkaline lysis maxiprep ^34, 35^. Competent cells were transformed with plasmids, plated to obtain single colonies, and inoculated into LB medium with appropriate antibiotics. Primary cultures were grown for 14 h at 37°C. Cells were pelleted (6,000 × g, 10 min, 4°C), resuspended in Solution I (25 mM Tris-HCl pH 8.0, 10 mM EDTA), and lysed with Solution II (0.2 N NaOH, 1% SDS) for 10 min at room temperature with gentle mixing. Solution III (3 M potassium acetate, 2 M glacial acetic acid) was added, mixed gently, and incubated on ice for 20 min. Lysates were centrifuged (14,000 × g, 20 min, 4°C), and supernatants were precipitated with isopropanol (20–30 min, room temperature). Pellets were washed with 70% ethanol, dried, treated with RNase A (37°C), and extracted with phenol:chloroform followed by chloroform. DNA was precipitated with isopropanol and 3 M sodium acetate, pelleted (10,000 × g, 10 min), washed with 70% ethanol, and resuspended in 1× TE buffer (10 mM Tris-HCl, 1 mM EDTA). Plasmid quality was verified by 0.8% agarose gel electrophoresis.

### NHEJ reporter assay

Intracellular NHEJ activity was measured as described previously ^21, 32, 36, 37^. Briefly, Molt4 (4 × 10L cells) and Jurkat cells were seeded in 6-well plates and transfected with 20 μg I-SceI overexpression construct and 15 μg pimEJ5-GFP reporter using branched PEI, with coffee decoction at increasing concentrations (Molt4: 0.1–1.6 mg/mL; Jurkat: 0.1–0.5 mg/mL). Cells were incubated at 37°C for 48 h, harvested, and analysed by flow cytometry (CytoFLEX S, Beckman Coulter, USA) to quantify GFP-positive cells. Percent GFP+ cells were plotted as bar graphs.

### Immunofluorescence

Cells were seeded on coverslips (5 × 10L cells/mL) and cultured for 24 h, then treated with coffee decoction (0.1–3.2 mg/mL, 5 h, 37°C, 5% CO₂) for immunofluorescence studies as described ^38, 39^. Cells were washed with PBS, fixed in 2% paraformaldehyde (10 min, RT), permeabilized with PBST (0.1% Triton X-100, 10 min, RT), blocked with PBST + 0.1% BSA + 10% FBS (1 h, 4°C), and incubated overnight at 4°C with primary antibody [γ-H2AX (Cell Signaling Technology, USA)]. Alexa Fluor-conjugated secondary antibodies (Life Technologies, USA) were added (2 h, RT), followed by mounting in DAPI/DABCO. Images were acquired on a laser scanning confocal microscope (Olympus FV3000, Japan) and processed with Olympus software.

For time-dependent DSB resolution post-IR, HeLa cells were pretreated with 0.4 mg/mL coffee decoction (5 h), irradiated (5 Gy), and harvested at 0.5, 4, 18, or 24 h post-IR for immunofluorescence as above.

### Overexpression and purification of His-tagged Ligase IV/XRCC4

The His-tagged Ligase IV/XRCC4 co-expression plasmid was a gift from Dr. Mauro Modesti ^23^. *Rosetta*(DE3)pLysS cells were transformed, grown to OD₆₀₀ = 0.6, and induced with 1 mM IPTG (16 h, 16°C). Cells were harvested (6,000 × g), resuspended in extraction buffer [20 mM Tris-HCl pH 8.0, 0.5 M KCl, 20 mM imidazole pH 7.0, 20 mM β-mercaptoethanol, 10% glycerol, 0.2% Tween 20, 1 mM PMSF], and lysed. Clarified lysate was loaded onto Ni-NTA resin (Novagen, USA) per manufacturer’s instructions. Eluates were pooled, applied to a UNOsphere Q anion-exchange column (Bio-Rad, USA), and proteins were eluted with a KCl gradient. Peak fractions were dialyzed overnight into 20 mM Tris-HCl pH 8.0, 150 mM KCl, 2 mM DTT, 10% glycerol, snap-frozen, and stored at −80°C. Purity and identity were confirmed by SDS-PAGE ^23, 32, 40^.

### DNA ligation reactions using purified protein

DNA ligation reactions with purified Ligase IV/XRCC4 were performed to assess coffee decoction effects. Purified protein was preincubated with coffee decoction (0.625-40 mg/mL) in joining buffer [30 mM HEPES-KOH pH 7.9, 7.5 mM MgCl₂, 1 mM DTT, 2 mM ATP, 50 μM dNTPs, 0.1 μg BSA] for 30 min at 25°C ^41, 42^, followed by addition of radiolabelled compatible-end DNA substrate and further incubation (1 h, 25°C). Products were phenol:chloroform extracted, ethanol/glycogen precipitated, resuspended in 10 μL TE buffer, resolved on 8% denaturing PAGE, dried, and visualized using a phosphorImager (FLA9000, Fuji, Japan). Joined products were quantified in PSLU using MultiGauge software.

### Complementation Assay with Purified Ligase IV/XRCC4

Caffeine-mediated NHEJ suppression by coffee decoction (30 mg/mL) in cell-free extracts was restored by adding purified Ligase IV/XRCC4, followed by incubation (1 h, 25°C). Reactions were terminated, DNA purified by phenol:chloroform extraction and glycogen/ethanol precipitation, resuspended in 10 μL TE buffer, heat-denatured, and resolved by 8% denaturing PAGE. Joined products were quantified in PSL Units using MultiGauge software and plotted as bar graphs in GraphPad Prism.

### Statistical analysis

Statistical analysis was carried out using GraphPad Prism software. Significance was determined using either Students’ t-test or one-way ANOVA. The obtained values were considered significant if the p-value was less than 0.05 and *, **, ***, **** denotes p< 0.05, p< 0.005, p<0.001, and p<0.000, respectively.

## Results

### Determination of caffeine content in the Coffee powder decoction

Coffee powder was sourced from farms in Kerala, India. Samples containing 10, 20, 40, 60 or 80 mg were weighed, resuspended in 1 mL autoclaved double-distilled water, vortexed for 1 min, and heated at 95°C for 10 min. The mixtures were then centrifuged (5000 rpm, 10 min, room temperature), and the resulting supernatant (coffee decoction) was used for experiments after caffeine quantification.

Caffeine concentrations were determined by high-performance liquid chromatography (HPLC) with UV detection at 273 nm. A standard curve was generated using pure caffeine (Sigma) at 100, 250, 500, 750, and 1000 µg/mL (Figure S1). Retention time (*t_R_*) provided qualitative identification, while peak area enabled quantification (Figure S2A). Plotting peak area against concentration yielded a linear standard curve (*Y* = 14173.8*X* + 93041.1 *R*^2^ = 0.999; Figure S2B, C), confirming method suitability. The of pure caffeine was 4.9 min.

Coffee decoctions (10–100 mg/mL) were analyzed similarly (Figure 1A-C). Caffeine was identified by matching *t_R_* (4.9 min) and spectral profiles to the pure standard (Figure 1C-F, S1, S2A). The decoction-specific standard curve was also linear (Figure 1D, E). Notably, the peak area for 40 mg/mL decoction matched that of 750 µg/mL pure caffeine (Figure 1F), indicating equivalent caffeine content. The coffee beans powder (CBP) was then used for subsequent DNA repair studies.

**Figure 1.**
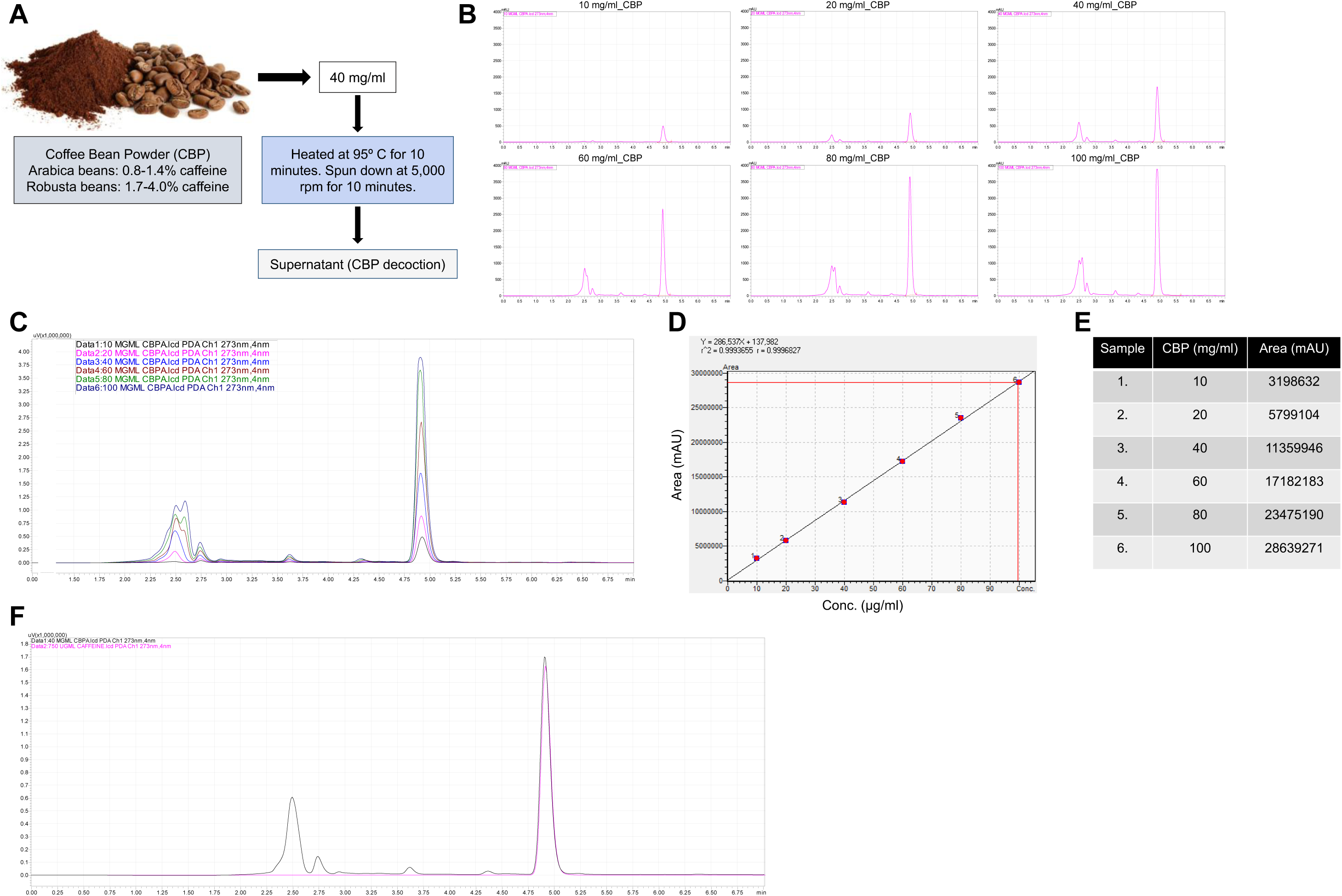
Estimation of caffeine content in coffee bean powder sample. **A.** Schematic representation of the steps followed to prepare coffee decoction from coffee bean powder. **B.** Histograms representing peaks of caffeine following HPLC studies when the increasing concentrations (10, 20, 40, 60, 80, and 100 mg/ml) of coffee decoction samples were used. **C.** Histograms representing the overlay of peaks of caffeine in the increasing concentrations (10, 20, 40, 60, 80 and 100 mg/ml) of coffee decoction prepared from coffee bean powder after HPLC studies. **D.** Graph representing the area under the peak of caffeine when the increasing concentrations (10, 20, 40, 60, 80, and 100 mg/ml) of coffee decoction samples were subjected to HPLC studies. **E.** Table showing the area under the curve of the caffeine peak for each concentration of coffee decoction used for the HPLC analysis. **F.** Histogram from HPLC studies representing the overlay of the area under the peak of 750 µg/ml of pure caffeine (Sigma) to that of the 40 mg/ml of coffee decoction sample.

### Coffee decoction inhibits the end joining of DNA double-strand breaks

The effect of coffee decoction on NHEJ was investigated using the cell-free repair assay system. In order to assess the effect of coffee decoction on end joining of compatible ends DNA substrate, 75 nt long double-stranded oligomeric DNA substrate, SCR19/20 harbouring 5’-5’ compatible ends (Figure 2A, S3A) were incubated with increasing concentrations of coffee decoction (1.25, 2.5, 5, 10, 20 and 40 mg/mL) in the presence of Molt4 or Jurkat cell-free extracts (Figure 2B, C; S3B, S4A, B). The effect of coffee decoction on joined products formation was evaluated by resolving the purified DNA products on a denaturing polyacrylamide gel (Figure S3A). A concentration-dependent decrease in the efficiency of joined product formation was observed when increasing concentrations (1.25, 2.5, 5, 10, 20 and 40 mg/mL) of coffee decoction was incubated with cell extracts and DNA substrate. The highest level of inhibition was noted at the higher decoction concentrations of 10, 20 and 40 mg/ml (Figure 2B, C and S4A, B). A significant reduction in joining efficiency was observed at concentrations ≥5 mg/mL. In contrast, reactions containing 1.25 and 2.5 mg/mL of CBP decoction showed joining levels comparable to the protein-alone control, indicating minimal or no inhibitory effect at these lower concentrations (Figure 2B, C; S4A, B). This indicates that caffeine present in the coffee decoction was able to inhibit the end joining of compatible end DNA substrate.

**Figure 2.**
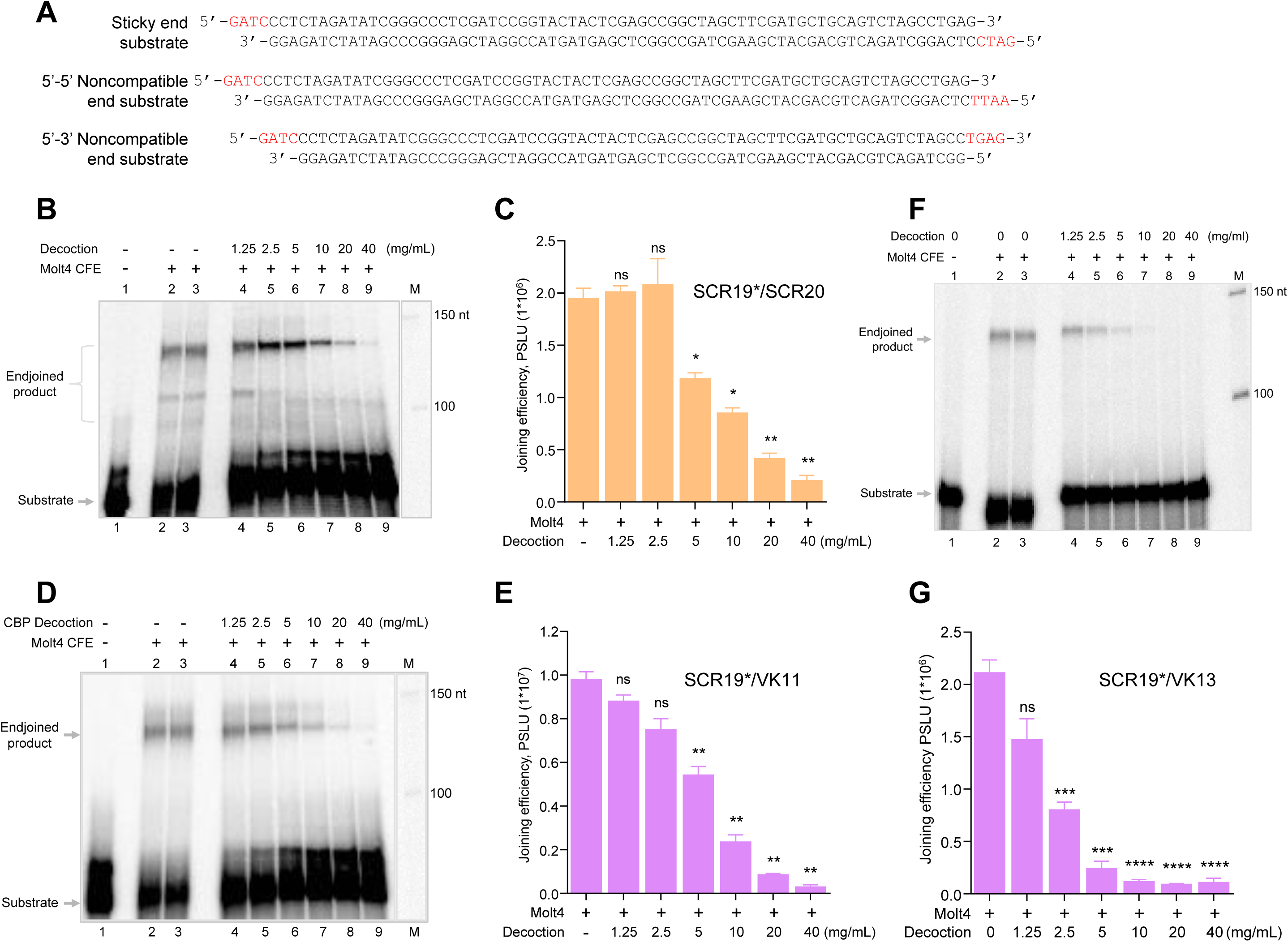
Evaluation of the impact of the coffee decoction on NHEJ mediated joining of compatible and noncompatible ends DNA substrates. **A.** Figure representing the sequence of compatible end DNA substrate (SCR19/20) and noncompatible ends (SCR19/VK11 and SCR19/VK13) substrate used for the joining assay. **B.** Representative denaturing PAGE showing the effect of the increasing concentration of coffee decoction on the joining of compatible end substrate (SCR19/20) when the DNA was incubated with Molt4 cell-free extract. **C.** The experiment shown in panel B was repeated three times, and the joined product was quantified using Multi Gauge V3.0 software and presented as a bar graph showing mean ± SEM. **D.** Representative image of the denaturing PAGE depicting the impact of coffee decoction on the end joining of 5’-5’ noncompatible ends DNA substrate (SCR19/VK11) in the presence of Molt4 CFE. **E.** The bar diagram showing the quantitation of the experiment showed in panel D (n=3); the data represented is mean ± SEM. **F.** Representative denaturing PAGE illustrating the impact of increasing concentration of coffee decoction on the joining 5’-3’ noncompatible end substrate (SCR19/VK13) when the DNA was incubated with Molt4 cell-free extract. **G.** The experiment shown in panel F was repeated three times, and the joined product was quantified using Multi Gauge V3.0 software and presented as a bar graph showing mean ± SEM. (ns: not significant, *p<0.05, **p<0.005, ***p<0.001, ****p<0.0001).

To evaluate the effect of coffee decoction on end joining of DSBs with 5’-5’ noncompatible ends DNA substrate, radiolabelled double-stranded oligomeric DNA substrate harbouring 5’-5’ noncompatible ends (SCR19/VK11) (Figure 2A) were incubated with increasing concentrations of coffee decoction (1.25, 2.5, 5, 10, 20 and 40 mg/mL) in the presence of Molt4 or Jurkat CFE (Figure 2D-E; S5A-D). It was observed that the formation of joined product was inhibited at coffee decoction concentrations of ≥5 mg/mL when incubated with Molt4 or Jurkat CFE (Figure 2D, E; S5A-D). In the case of noncompatible ends substrate (SCR19/VK11), we observed an increase in the joining efficiency at lower concentrations of coffee decoction (1.25 and 2.5 mg/ml), however this observed increase in joining efficiency was significant only in case of Molt4 CFE and not in case of Jurkat CFE (Figure S5B, D). It is possible that lower concentrations of coffee decoction inhibit nuclease activity present in the cell-free extracts, thereby reducing end resection and resulting in enhanced joining efficiency at these concentrations. However, this possibility warrants further experimental validation. Overall, these findings demonstrate that coffee decoction inhibits the joining of noncompatible DNA ends, thereby indicating its capacity to suppress NHEJ activity. When 5’-3’ noncompatible ends (SCR19/VK13) were used, a concentration-dependent inhibition in the joining was observed in the case of Molt4 and Jurkat cell extracts in the presence of coffee decoction (1.25, 2.5, 5, 10, 20 and 40 mg/mL) (Figure 2F, G; S4C, D). >50% of the inhibition of NHEJ was observed when incubated with 5 mg/mL of coffee decoction in both the cell lines. The effect was more pronounced when the the 5’-3’ noncompatible ends DNA substrate (SCR19/VK13) was incubated with a higher concentration of coffee decoction (10, 20 and 40 mg/mL) in the presence of Jurkat as well as Molt4 cell-free extract (Figure 2F, G; S4C, D).

Highly proliferative cancer cells were employed for these studies, as they experience elevated replication stress and rely heavily on DNA double-strand break (DSB) repair pathways to preserve genomic stability and sustain rapid proliferation. Consequently, they provide a sensitive and biologically relevant model to evaluate the impact of coffee decoction on NHEJ activity. To determine whether coffee decoction-mediated inhibition of end joining was specific to cancer cells, we next examined NHEJ efficiency in normal yet transformed fibroblast cells (HMF-3S). Incubation of oligomeric DNA substrates bearing compatible or noncompatible ends with HMF-3S cell-free extracts (CFE) in the presence of coffee decoction (1.25, 2.5, 5, 10, 20 and 40 mg/mL) resulted in a concentration-dependent reduction in joining efficiency (Figure 3A-F). A pronounced decrease in end-joining activity was observed at concentrations ≥5 mg/mL of coffee decoction.

**Figure 3.**
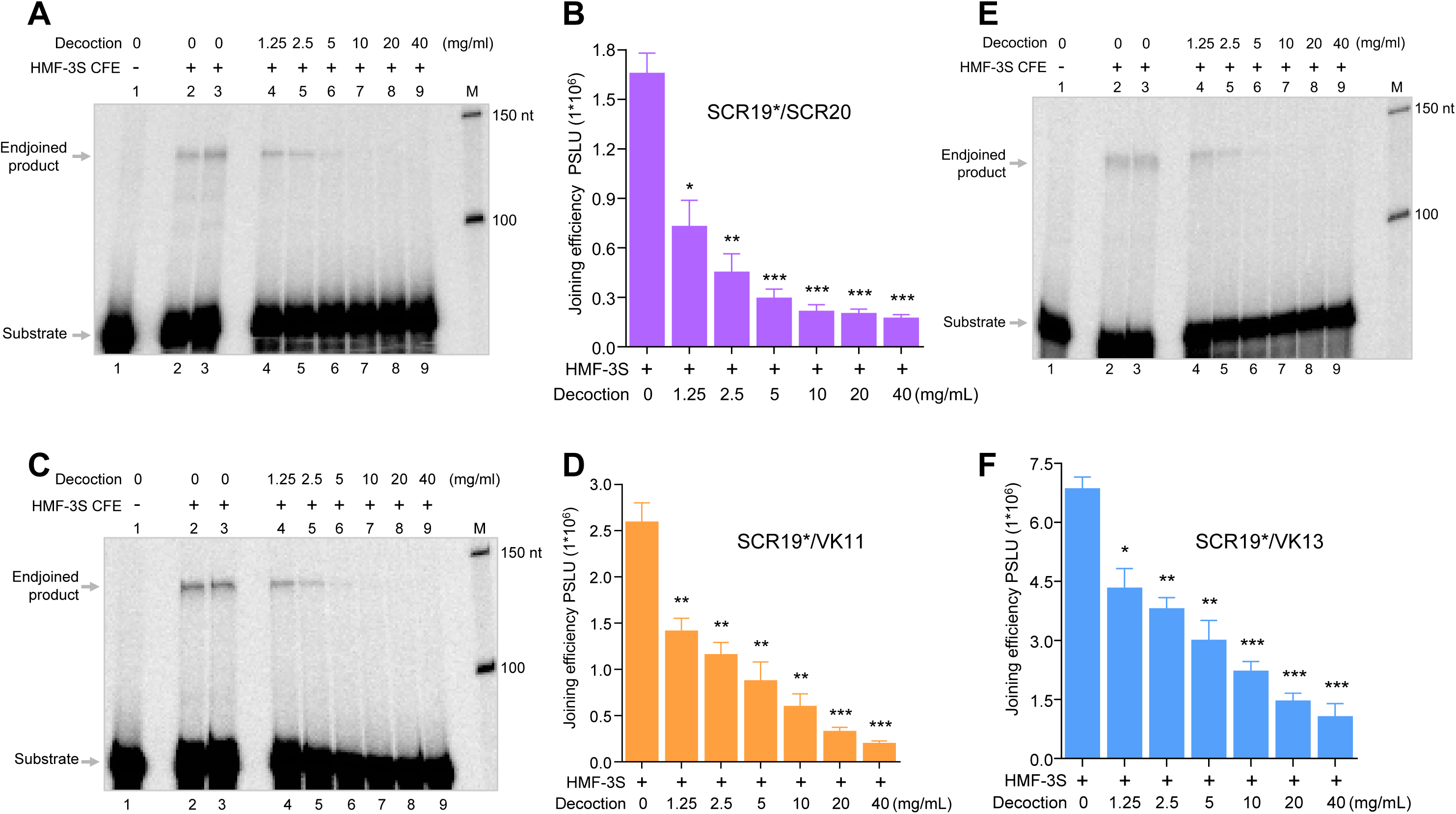
Evaluation of the impact of the coffee decoction on NHEJ mediated joining of compatible and noncompatible ends DNA substrates in the presence of cell-free extract derived from primary cell line. **A.** Representative denaturing PAGE showing the effect of the increasing concentration of coffee decoction on the joining compatible end substrate (SCR19/20) when the DNA was incubated with HMF-3S cell-free extract. **B.** The experiment shown in panel A was repeated three independent times, and the joined product was quantified using Multi Gauge V3.0 software and presented as a bar graph showing mean ± SEM. **C.** Denaturing PAGE profile depicting the impact of increasing concentrations of coffee decoction on the end joining of 5’-5’ noncompatible ends DNA (SCR19/VK11) in the presence of HMF-3S CFE. **D.** The experiment shown in panel C was repeated three times, and the joined product was quantified using Multi Gauge V3.0 software and presented as a bar graph showing mean ± SEM. **E.** Denaturing PAGE profile depicting the concentration dependent effect of coffee decoction on the end joining of 5’-3’ noncompatible ends DNA (SCR19/VK13) in the presence of HMF-3S CFE. **F.** The experiment shown in panel E was repeated thrice, and the joined product was quantified using Multi Gauge V3.0 software and presented as a bar graph showing mean ± SEM. (ns: not significant, *p<0.05, **p<0.005, ***p<0.001, ****p<0.0001).

### Addition of coffee decoction leads to reduced NHEJ activity within the human cells

An extrachromosomal assay system was used to evaluate the effect of coffee decoction on NHEJ within human cells, as previously described ^20, 21, 32, 36^ (Figure 4A, B). The NHEJ reporter construct pimEJ5GFP, with a disrupted GFP gene flanked by I-SceI sites ^21, 32, 36^, was employed. I-SceI-induced DSBs in transfected pimEJ5-GFP (pJS296) episomes are repaired via NHEJ, restoring GFP expression. Molt-4 cells were transiently co-transfected with pimEJ5GFP and I-SceI overexpression constructs along with increasing decoction concentrations (0.1–1.6 mg/mL; Figure 4C, D). After 48 h, flow cytometry measured GFP-positive cells as a readout of NHEJ repair (Figure 4C, D). Results showed that the coffee decoction reduced GFP-positive cells in a concentration-dependent manner, indicating intracellular NHEJ inhibition even within the human cells (Figure 4C, D).

**Figure 4.**
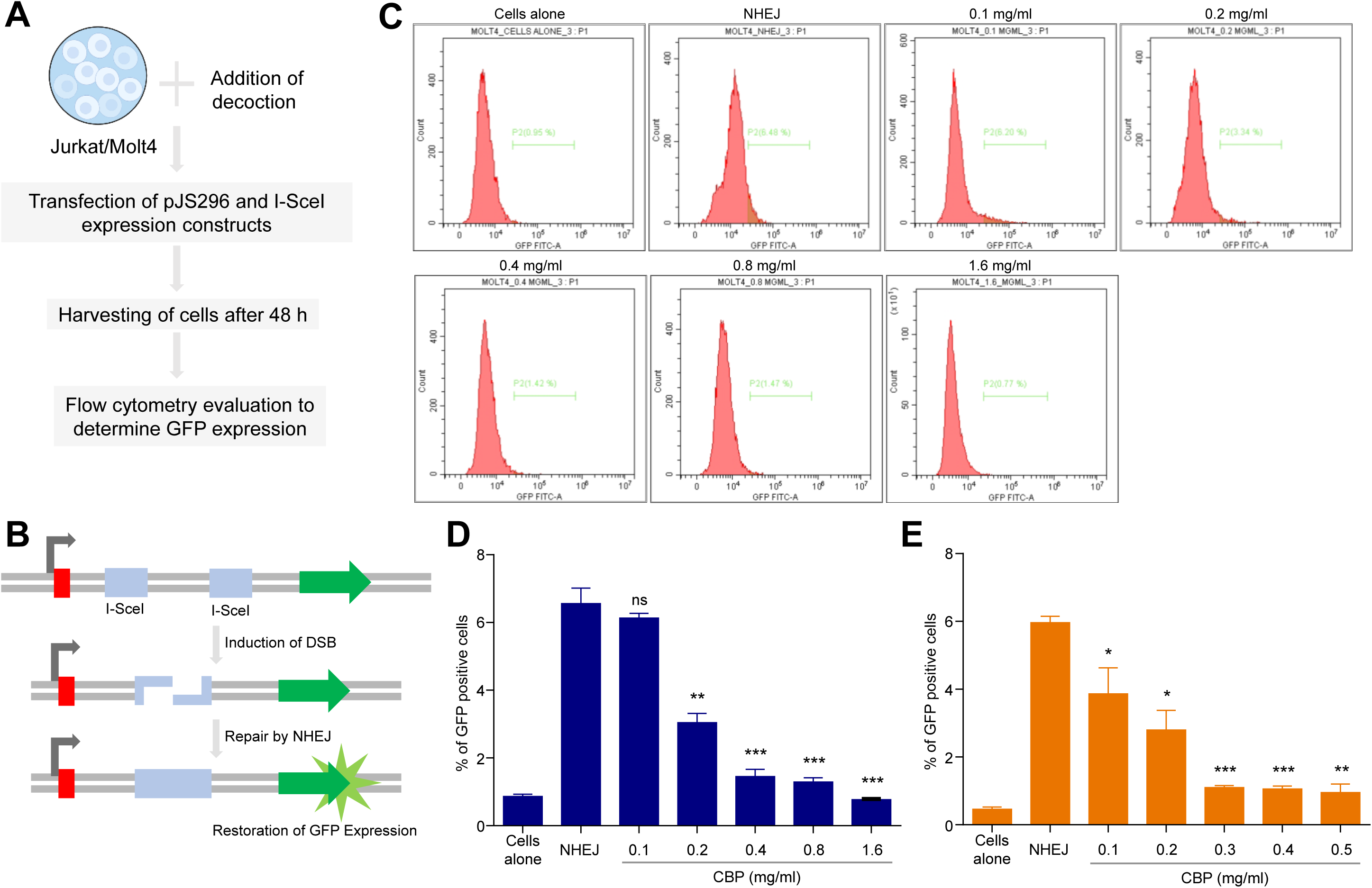
Impact of coffee decoction on extrachromosomal NHEJ in Molt4 and Jurkat cells. **A.** Representative image illustrating the steps being followed to assess the impact of caffeine on NHEJ in human cells. **B.** Schematic representation of the intracellular NHEJ assay. The PimEJ5GFP reporter construct contains a disrupted GFP sequence flanked by I-SceI recognition sites. Transfection with an I-SceI overexpression construct induces a double-strand break in PimEJ5GFP. Successful repair through NHEJ restores GFP expression, serving as a readout for NHEJ efficiency. **C.** Histograms representing the comparison of intracellular NHEJ activity in Molt4 cells in presence of increasing concentrations of coffee decoction as compared to untreated cells. **D.** GFP positive cells, post-transfection of Molt4 cells with PimEJ5GFP and I-SceI overexpression constructs in the presence of increasing concentrations (0.1, 0.2, 0.4, 0.8, and 1.6 mg/ml) of coffee decoction were analyzed by flow cytometry and plotted as bar graph representing mean ± SEM. **E.** Bar graph represent the percentage of GFP-positive cells as a measure of intracellular NHEJ activity upon transfection of PimEJ5GFP and I-SceI overexpression constructs in Jurkat cells in the presence of coffee decoction (0.1, 0.2, 0.3, 0.4 and 0.5 mg/ml). The error bar represents mean ± SEM. (ns: not significant, *p<0.05, **p<0.005, ***p<0.001, ****p<0.0001).

The experiment was replicated in Jurkat cells, showing significant decreases in GFP-positive cells with rising decoction concentrations (Figure 4E). These results confirm that caffeine in coffee decoction inhibits NHEJ in human cells, which was consistent with the effect of purified caffeine ^20^.

### Inhibition of NHEJ at chromosomal level by coffee decoction leads to the accumulation of unrepaired DSBs in human cells

Based on the above observations, we next investigated whether inhibition of NHEJ by coffee decoction leads to the accumulation of unrepaired endogenous chromosomal DSBs at the genome-wide level. To address this, HeLa cells were treated with increasing concentrations of coffee decoction, followed by immunofluorescence analysis using a γ-H2AX antibody (Figure 5A). γ-H2AX foci were used as a surrogate marker of DSBs to assess the impact of coffee decoction on chromosomal DNA damage. We observed a clear increase in γ-H2AX foci in HeLa cells with rising concentrations of coffee decoction (0.1, 0.4, 1.6, and 3.2 mg/mL), indicating a dose-dependent accumulation of unrepaired DSBs (Figure 5B, C, S6A, B). This increase in γ-H2AX signal suggests that caffeine present in the coffee decoction impairs NHEJ-mediated repair of chromosomal breaks. Consistent with this interpretation, treatment with SCR7 (50 µM), a known DNA ligase IV inhibitor, which was used as a positive control resulted in similar elevated γ-H2AX foci.

**Figure 5.**
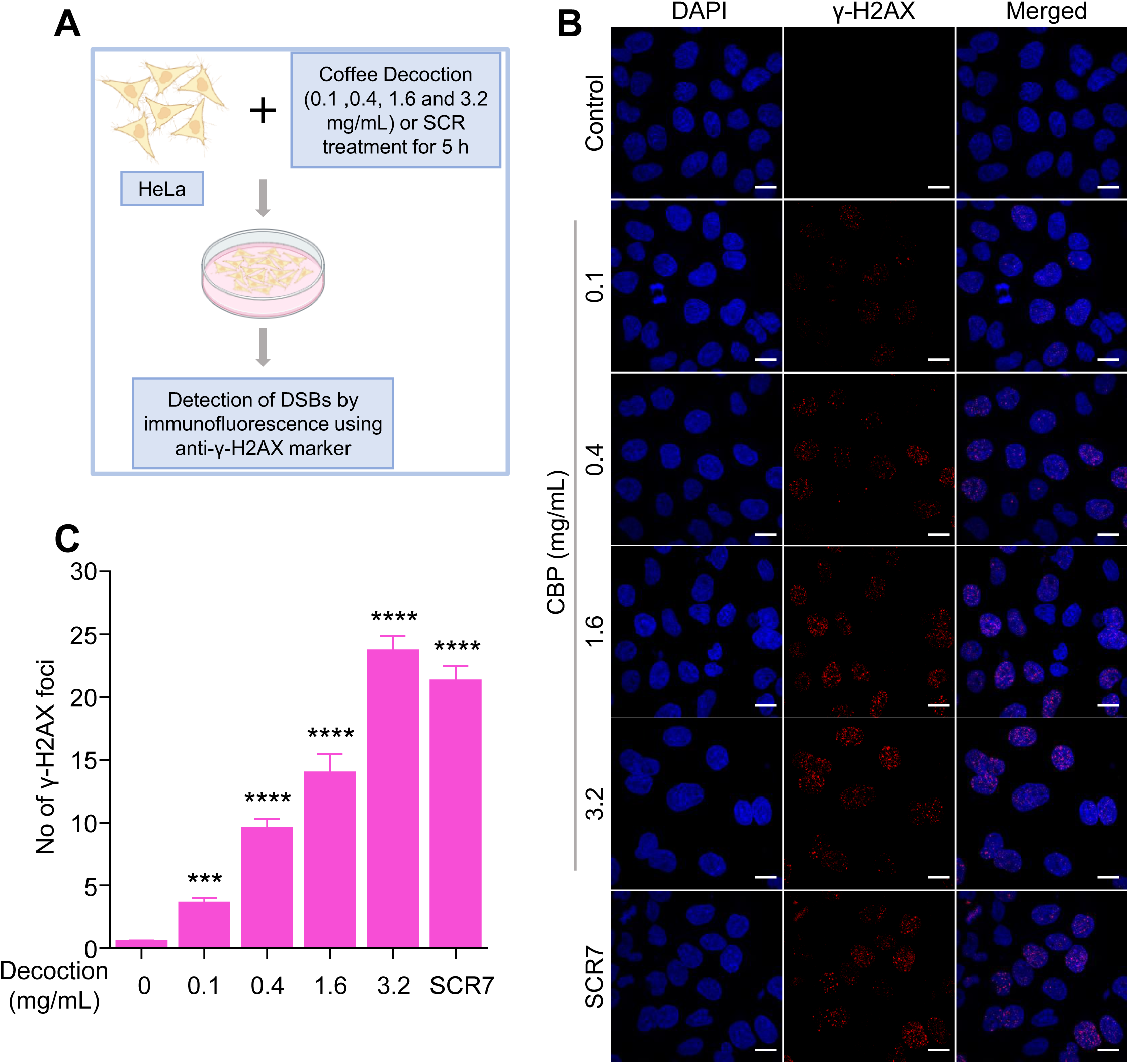
Evaluation of effect of coffee decoction on chromosomal DSB repair within the human cells. **A.** Schematic presentation depicting the steps followed while evaluating the impact of coffee decoction on the repair of DNA double-strand breaks within HeLa cells. **B.** Representative immunofluorescence images showing the impact of coffee decoction on DSB repair using γ-H2AX foci as a marker for DSBs inside nucleus. HeLa cells were treated with increasing concentrations (0.1, 0.4, 1.6, and 3.2 mg/ml) of coffee decoction for 5 h, following which the cells were proceeded for immunofluorescence and probed with γ-H2AX antibody. The nucleus was stained using DAPI. In each case, a merged image for both signals had been shown as a panel on the right. SCR7 (50 µM) was used as a positive control for NHEJ inhibition. Scale bar represents 10 µm. **C.** The experiment was repeated three times, and approximately 100 cells from each batch were evaluated for each sample. The average number of foci is represented as a bar graph showing mean ± SEM. (ns: not significant, *p<0.05, **p<0.005, ***p<0.001, ****p<0.0001).

Together, these findings indicate that caffeine in coffee decoction not only inhibits NHEJ in biochemical- or episome-based reporter systems but also effectively blocks NHEJ-mediated repair at the chromosomal level in mammalian cells, leading to the persistence and accumulation of unrepaired DSBs.

### Exposure to coffee decoction impairs DSB repair kinetics, leading to prolonged persistence of DNA damage

Building on our observation that coffee decoction causes accumulation of endogenous unrepaired chromosomal DSBs, we investigated whether this stems from defective DSB repair kinetics after genotoxic insult, rather than solely increased DSB induction. To test this, we monitored γ-H2AX foci dynamics in HeLa cells following exposure to ionizing radiation (IR; 5 Gy) with or without coffee decoction (0.4 mg/mL), a concentration chosen to mimic physiologically relevant exposure levels while minimizing overt cytotoxicity. γ-H2AX foci were rapidly induced 0.5 h post-IR in all groups, confirming efficient DSB formation across conditions (Figure 6A, B). In untreated controls, foci levels progressively declined by 12–24 h, reflecting robust NHEJ-mediated repair (Figure 6A, B). In contrast, coffee decoction-treated cells exhibited markedly prolonged γ-H2AX persistence at these time points, indicating substantially delayed DSB resolution (Figure 6A, B).

**Figure 6.**
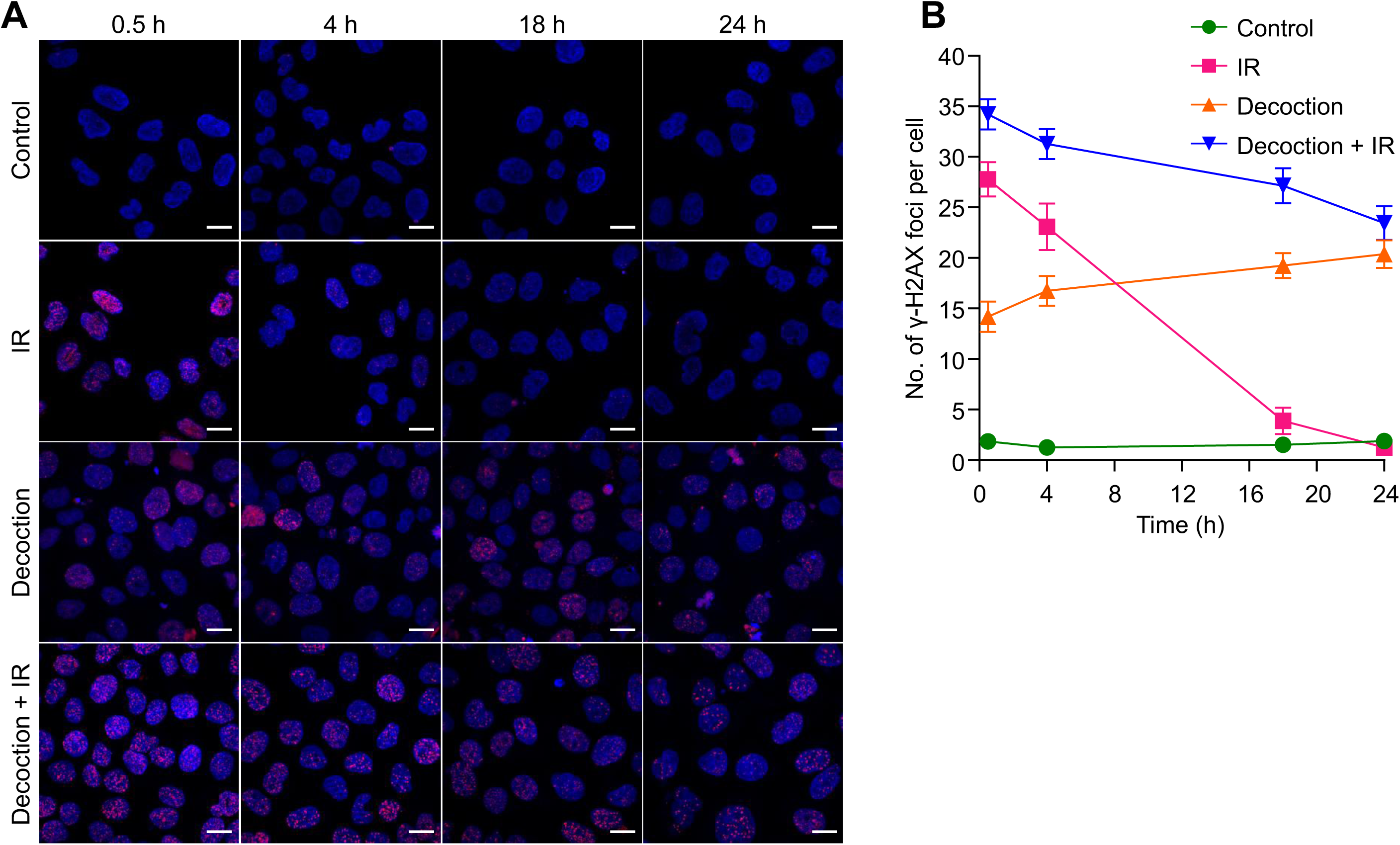
Effect of caffeine on DNA damage response in HeLa cells. **A.** Representative confocal images showing γ-H2AX foci (red) and nuclear DNA stained with DAPI (blue) in HeLa cells treated with IR (5 Gy) with or without coffee decoction treatment (0.4 mg/mL) at various time points (0.5, 4, 18 and 24 h) post-irradiation. Scale bar, 10 μm. **B.** Quantification of γ-H2AX foci in HeLa cells exposed to ionizing radiation (IR, 5 Gy) in the presence or absence of caffeine (0.4 mg/mL coffee decoction). Experiments were performed two independent times and cells were fixed at the indicated time points post-IR and immunostained for γ-H2AX. The average number of γ-H2AX foci per nucleus was quantified and presented as line plot showing mean ± SEM.

Notably, coffee decoction alone (without IR) triggered a significant elevation in baseline γ-H2AX foci relative to untreated cells, suggesting that caffeine induces DSBs independently, possibly through replication stress, topoisomerase inhibition or due to endogenous inhibition of the NHEJ (Figure 6A, B). When combined with IR, decoction further exacerbated foci retention compared to IR alone, underscoring impaired repair kinetics. Collectively, these data demonstrate that caffeine in coffee decoction not only generates DSBs but also compromises their timely repair, leading to sustained DNA damage signalling and potential genomic instability. This dual mechanism, DSB induction coupled with repair blockade, highlights coffee decoction’s potential as a radio sensitizing agent.

### Coffee decoction inhibits Ligase IV/XRCC4-mediated end joining, and purified Ligase IV/XRCC4 complementation reverses the inhibition

In a recent study, we demonstrated that caffeine inhibits nonhomologous end joining (NHEJ) by functionally compromising the Ligase IV/XRCC4 complex ^20^, establishing this core ligation machinery as caffeine’s direct molecular target. Building on these findings and our observations that coffee decoction suppresses NHEJ activity while delaying DSB repair kinetics, we hypothesized that decoction’s inhibitory effects similarly arise from direct interference with Ligase IV/XRCC4, the indispensable final step in classical NHEJ. Impairment here would leave persistent DNA ends and unrepaired breaks, phenocopying known mechanism. We thus tested whether coffee decoction directly disrupts Ligase IV/XRCC4-mediated ligation and addition of exogenous purified complex could rescue NHEJ in decoction-treated extracts.

To directly assess this, Ligase IV/XRCC4 was purified via Ni-NTA chromatography using an imidazole gradient, as described previously ^32^, with protein purity assessed by PAGE (Figure S7). Joining assays employed DSBs bearing compatible end DNA substrate (SCR19/20) incubated with purified Ligase IV/XRCC4 in presence of escalating decoction concentrations (0.625–40 mg/mL). Products were resolved by denaturing PAGE, revealing dose-dependent inhibition of joining (Figure 7A, B). These data confirm that caffeine in coffee decoction directly impairs Ligase IV/XRCC4 mediated joining of broken DNA.

**Figure 7.**
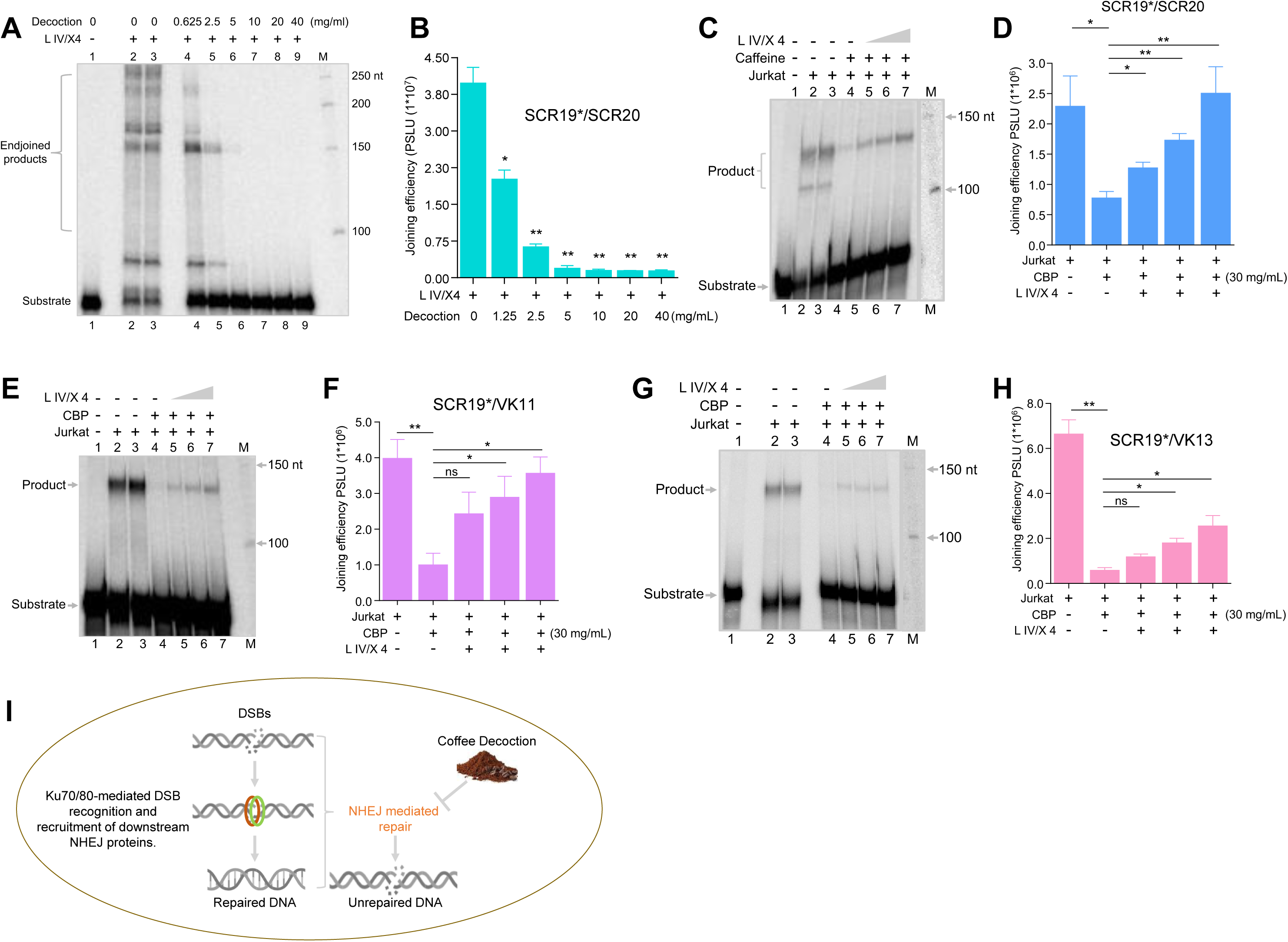
Evaluation of the effect of coffee decoction on joining of sticky DNA ends and reconstitution studies following coffee decoction-mediated inhibition of NHEJ. **A.** Representative denaturing PAGE gel profile showing the effect of coffee decoction on the ligation of compatible end substrate by purified Ligase IV/XRCC4 protein. **B.** The experiment shown in panel A was repeated three times, the joined product was quantified and presented as a bar graph, showing mean ± SEM. **C.** A representative denaturing PAGE showing reconstitution of the end joining of compatible end substrate (SCR19/20) catalyzed by Jurkat cell-free extract after inhibition due to coffee decoction treatment (30 mg/mL). Following caffeine treatment, Ligase IV/XRCC4 protein was added to determine, if the joining could be reconstituted. Initially, inhibition of end joining with cell-free extract in the presence of coffee decoction was conducted. Following the inhibition reaction, the increasing concentration of overexpressed and purified Ligase IV/XRCC4 was added back to the samples incubated with coffee decoction to evaluate the effect of caffeine on Ligase IV/XRCC4 mediated joining. **D.** The joined product was quantified and presented as a bar graph representing mean ± SEM (n=3). **E-H.** Representative denaturing PAGE image for end joining of 5’-5’ (SCR19/VK11, E) and 5’-3’ noncompatible end substrate (SCR19/VK13, G), respectively post Ligase IV/XRCC4 complementation upon treatment with coffee decoction in the presence of Jurkat cell-free extract. The joined products for the substrates SCR19/VK11 (F) and SCR19/VK13 (H) were quantified and presented as a bar graph. The experiments were conducted three independent times, and the error bar represents mean ± SEM. (ns: not significant, *p<0.05, **p<0.005, ***p<0.001, ****p<0.0001). **I.** Schematic of NHEJ-Mediated DSB Repair and Caffeine Inhibition. DNA double-strand breaks are primarily repaired via the nonhomologous end joining (NHEJ) pathway. The Ku70/Ku80 heterodimer rapidly binds DSB ends, recruiting core NHEJ factors (e.g., DNA-PKcs, Artemis, XRCC4-Ligase IV) for end processing and ligation, restoring genomic integrity. In the presence of caffeine from coffee decoction, NHEJ is inhibited; caffeine disrupts XRCC4-Ligase IV-mediated joining, impairing repair complex assembly and leading to persistent unrepaired DSBs within the cells.

A reconstitution assay further validated target specificity in human cell-free extracts. Jurkat CFE was preincubated with decoction (30 mg/mL) for 1 h to abrogate joining, followed by addition of purified Ligase IV/XRCC4 and compatible-end substrate (SCR19/20). Decoction alone abolished product formation (Figure 7C, D). Strikingly, exogenous addition of Ligase IV/XRCC4 partially restored joining in decoction-treated CFE, achieving levels approaching untreated controls (Figure 7C, D). This rescue demonstrates that caffeine’s NHEJ inhibition is Ligase IV/XRCC4-dependent, with the complex acting as the primary bottleneck.

Since most physiological DSBs bear noncompatible ends requiring end-processing before ligation, we extended rescue experiments to two distinct substrates: 5’-5’ (SCR19/VK11) and 5’-3’ overhangs (SCR19/VK13). Remarkably, purified Ligase IV/XRCC4 robustly restored joining for both, yielding substantial product increases despite decoction pretreatment (Figure 7E–H). This broad rescue across end configurations underscores caffeine’s selective targeting of ligation over upstream processing.

Collectively, these findings mechanistically link coffee decoction’s NHEJ suppression to direct Ligase IV/XRCC4 inhibition, mirroring purified caffeine. By blocking this essential step, decoction promotes unrepaired DSB persistence, positioning it and dietary caffeine, as a natural modulator of genomic repair fidelity with potential therapeutic implications for replication-stressed malignancies.

## Discussion

Coffee and tea consumption forms an integral part of daily life for millions worldwide, with coffee serving as the predominant source of caffeine intake across food products, beverages, and pharmaceuticals. While caffeine’s psychoactive and stimulating effects are well established, its influence on genomic integrity remains an active area of research. Our study provides novel evidence that caffeine derived directly from *Coffea arabica* powder decoction potently inhibits nonhomologous end joining (NHEJ), a primary DNA double-strand break (DSB) repair pathway, in a concentration-dependent manner, in mammalian cells (Figure 7I).

### Coffee decoction efficiently suppresses NHEJ by targeting Ligase IV/XRCC4

HPLC analysis quantified caffeine content in 40 mg/mL Kerala-sourced coffee bean powder decoction (CBP) as equivalent to 750 μg/mL purified caffeine (Sigma), enabling precise concentration-matched comparisons. In mammalian cell-free extracts, coffee decoction dose-dependently inhibited end joining of both compatible and noncompatible DNA ends, with maximal suppression at highest concentrations. Extrachromosomal NHEJ reporter assays in Jurkat and Molt4 cells confirmed intracellular inhibition, evidenced by reduced GFP-positive cells proportional to decoction concentration. This suppression triggered chromosomal DSB accumulation, as demonstrated by elevated γ-H2AX foci formation. Kinetic analysis post-ionizing radiation (5 Gy) in HeLa cells revealed significantly delayed γ-H2AX foci resolution in decoction-treated cells, confirming impaired endogenous NHEJ dynamics.

Purified Ligase IV/XRCC4 ligation assays established direct, dose-dependent inhibition of the catalytic ligation step by coffee decoction. Complementation experiments established the specificity, particularly the NHEJ activity in cell extracts was inhibited by coffee decoction was partially restored by adding purified Ligase IV/XRCC4 complex. These biochemical reconstitution studies identify this complex as caffeine’s primary molecular target, explaining the observed suppression across *in vitro*, extrachromosomal, and chromosomal NHEJ assays.

### Genomic Stability and Therapeutic Considerations

DSBs represent among the most lethal genomic lesions, with accurate NHEJ repair essential for maintaining cellular homeostasis, particularly in non-replicating cells where NHEJ predominates. Caffeine-mediated NHEJ inhibition could compromise genomic stability in high-consumption individuals, potentially promoting chromosomal aberrations, mutations, and apoptosis through unrepaired DSB persistence. Notably, low decoction concentrations paradoxically enhanced end-joining efficiency, revealing biphasic regulation that may involve differential effects on repair factors or cell cycle modulation. Further studies are needed to elucidate these concentration-dependent mechanisms.

Our findings carry dual clinical implications. In cancer therapy, caffeine’s NHEJ suppression could sensitize tumors to DNA-damaging radiotherapy and chemotherapy by preventing repair of treatment-induced DSBs, building on its established radiosensitizing properties. Conversely, high dietary intake poses genotoxic risks to normal cells, warranting caution in therapeutic applications. With caffeinated products consumed by ∼80% of the global population daily, these results necessitate re-evaluation of caffeine’s long-term impact on population-level DNA repair capacity.

### Broader Public Health Perspective

Moderate caffeine consumption offers neuroprotection and reduces disease risk, yet higher doses impair critical DNA repair processes, warranting caution. As a ubiquitous dietary staple, coffee consumption modulates DSB repair efficiency, especially among heavy users, posing risks to genomic maintenance while opening avenues for repurposing caffeine as an NHEJ-targeting adjuvant in precision cancer therapeutics.

At low to moderate concentrations (0.1–1 mM, achievable via typical intake), caffeine exerts radioprotective effects by scavenging ROS from ionizing radiation (IR), thereby curbing oxidative DNA damage, lipid peroxidation, chromosomal aberrations, and micronucleus formation while preserving cellular redox balance, DNA integrity, and normal cell cycle progression ^43, 44, 45^. In contrast, higher doses produce a dual inhibitory effect: beyond suppressing ATM/ATR kinase activity and checkpoint signaling, caffeine directly disrupts NHEJ complex assembly by interfering with the Ligase IV/XRCC4 interaction. This hinders XRCC4 recruitment to damage sites, compromising the terminal ligation step and overall repair fidelity HR ^46^.

In conclusion, this study provides compelling evidence that caffeine, a common dietary component, significantly affects the DNA damage response by impairing NHEJ-mediated repair. The widespread consumption of coffee and other caffeinated products underscores the need for deeper investigation into the long-term consequences of caffeine on maintenance of genome integrity. While the genotoxic effects observed at the higher concentrations of coffee decoction used raise concerns, they also highlight potential opportunities for leveraging caffeine’s properties in cancer therapy.

## Conflict of interest

The authors declare that they have no conflicts of interest with the contents of this article.

## Author contributions

Conceptualization, SCR; Design of Experiments, SCR, SK; Investigation, SK; Writing, SK, SCR; Funding Acquisition, SCR; Resources and Supervision, SCR.

## Data Availability

All data relevant to the article are presented as main figures and supplementary figures. Any additional information related to the data reported in this article will be shared by the lead contact upon request.

## Acknowledgments

We thank Dr Shivangi Sharma and members of the SCR laboratory for critical reading and comments on the manuscript. We also thank Dr Bibha Choudhary and Dr Varsha KK for her help. We also acknowledge the Central Animal Facility of IISc. This work was supported by grants from DAE, India (Grant No. 21/01/2016-BRNS/35074) and ICMR, India (Grant No. 5/1327/SCR/ICRC/2022/NCD-III) to SCR and DBT-Glue Grant (BT/PR23078/MED/29/1253/2017) to SCR. DST-FIST (SR/FST/LSII-045/2016) is also acknowledged for its support in maintaining the Central Facility of Biochemistry, IISc. SCR is a JC Bose National Fellow (JCB/2023/000041). SK is supported by Senior Research Fellowship from IISc.

## Notes

### Competing Interest Statement

The authors have declared no competing interest.

