## Supplementary Text for "High Consumption of Coffee Disrupts Nonhomologous End Joining Implications for Genomic Stability"

**Supplementary Figure Legends**

**Table S1. Oligomers used in the study.**

**Figure S1.** **HPLC analysis of commercially available caffeine. A.** Histograms representing peaks for the increasing concentration (100, 250, 500, 750 and 1000 µg/ml) of pure caffeine (Sigma).

**Figure S2. HPLC-based determination of the standard curve for pure caffeine. A.** Histogram representing overlay of area under peak of increasing concentration (100 (green), 250 (red), 500 (blue), 750 (pink) and 1000 (black) µg/ml) of pure caffeine (Sigma). **B.** Standard curve of caffeine prepared by plotting the concentration against the area under the peak. **C.** Table representing the area under the curve of the caffeine peak for each concentration of caffeine used for the HPLC analysis.

**Figure S3. Schematic representation of the strategy for assessing the impact of coffee decoction on NHEJ catalysed by cell-free extracts. A.** The cell-free extract was incubated with a 5′-end-labeled DNA substrate containing DSBs in the presence of increasing concentrations of coffee decoction. The reaction was terminated using Tris-EDTA, and the joined products were purified by phenol:chloroform extraction, precipitated with glycogen and ethanol, resuspended in TE, and analyzed on a 8% denaturing PAGE. "M" represents the marker, "S" denotes the substrate, and "P" indicates the joined products. **B.** Representative SDS-PAGE analysis of cell-free extracts from Molt4, Jurkat, and HMF-3S cell lines. Equal amounts of total protein from each cell line were resolved on SDS-PAGE and stained with Coomassie Brilliant Blue to visualize protein profiles.

**Figure S4. Effect of coffee decoction on end-joining activity catalysed by Jurkat cell-free extracts. A.** Representative denaturing PAGE analysis showing the effect of coffee decoction on the joining of compatible DNA end substrate (SCR19/20) following incubation with Jurkat cell-free extract. **B.** Bar graph representing the quantification of the experiment shown in panel A (n=3), with data presented as mean ± SEM. **C.** Denaturing PAGE profile showing the concentration dependent effect of coffee decoction on the joining 5’-3’ noncompatible end substrate (SCR19/VK13) when the DNA was incubated with Jurkat cell-free extract. **D.** The experiment shown in panel C was repeated three independent times, and the joined product was quantified using Multi Gauge V3.0 software and presented as a bar graph showing mean ± SEM. (ns: not significant, *p<0.05, **p<0.005, ***p<0.001, ****p<0.0001).

**Figure S5. The impact of addition of coffee decoction on NHEJ when noncompatible DNA ends were used. A.** Denaturing PAGE profile showing the effect of increasing concentrations of coffee decoction (0.16, 0.32, 0.63, 1.25, 2.5, 5, 10, 20, 20 and 40 mg/mL) on the end joining of noncompatible ends (SCR19/VK11) DNA in the presence of Molt4 CFE. **B.** The experiment shown in panel A was repeated three independent times, and the joined product was quantified using Multi Gauge V3.0 software and presented as a bar graph showing mean ± SEM. **C.** Denaturing PAGE profile depicting the impact of increasing concentrations of coffee decoction on the end joining of noncompatible ends DNA (SCR19/VK11) in the presence of Jurkat CFE. **D.** Bar diagram showing the quantitation of the experiment depicted in panel C (n=3) showing mean ± SEM. (ns: not significant, *p<0.05, **p<0.005, ***p<0.001, ****p<0.0001).

**Figure S6. Evaluation of status of DSB repair following addition of coffee decoction in HeLa cells using γ-H2AX staining. A.** Representative immunofluorescence images illustrating the effect of coffee decoction on DSB accumulation, as assessed by γ-H2AX foci formation. HeLa cells were treated with increasing concentrations of coffee decoction (0.1, 0.4, 1.6, and 3.2 mg/mL) for 5 h, followed by immunofluorescence staining using an anti–γ-H2AX. Nuclei were counterstained with DAPI. Merged images displaying γ-H2AX and DAPI signals are shown in the right panel for each condition. **B.** The experiment was repeated three independent times, and ~100 cells from each biological repeat were evaluated. The average number of foci is represented as a bar graph showing mean ± SEM. (ns: not significant, *p<0.05, **p<0.005, ***p<0.001, ****p<0.0001).

**Figure S7. Overexpression and purification of mammalian Ligase IV/XRCC4.** SDS-PAGE profile showing the purified His-tagged Ligase IV/XRCC4 protein. ‘M” denotes marker and lanes 1 and 2 indicate different fractions.
