## Supplementary Figure for "High Consumption of Coffee Disrupts Nonhomologous End Joining Implications for Genomic Stability"

Table S1. Oligomers used for the study.

| Oligomer | Sequence |
|---|---|
| SCR19 | 5’-GATCCCTCTAGATATCGGGCCCTCGATCCGGTACTACTCGAGCCGGCTAGCTTCGATGCTGCAGTCTAGCCTGAG-3’ |
| SCR20 | 5′-GATCCTCAGGCTAGACTGCAGCATCGAAGCTAGCCGGCTCGAGTAGTACCGGATCGAGGGCCCGATATCTAGAGG-3’ |
| VK11 | 5´-AATTCTCAGGCTAGACTGCAGCATCGAAGCTAGCCGGCTCGAGTAGTACCGGATCGAGGGCCCGATATCTAGAGG-3´ |
| VK13 | 5´-GGCTAGACTGCAGCATCGAAGCTAGCCGGCTCGAGTAGTACCGGATCGAGGGCCCGATATCTAGAGG-3´ |

Figure S1

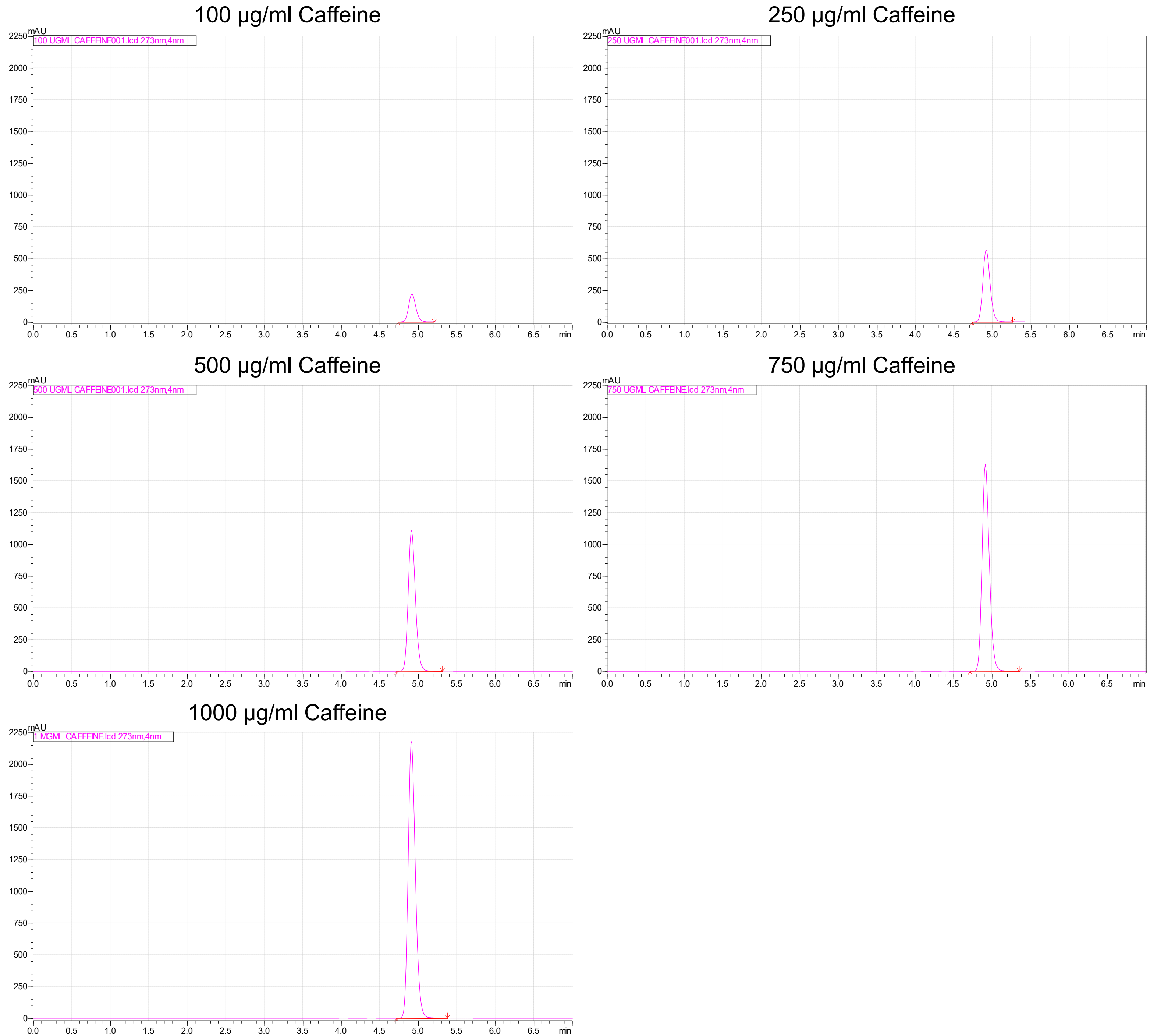
100 µg/ml Caffeine
100 UGML CAFFEINE001.lcd 273nm,4nm
250 µg/ml Caffeine
250 UGML CAFFEINE001.lcd 273nm,4nm
500 µg/ml Caffeine
500 UGML CAFFEINE001.lcd 273nm,4nm
750 µg/ml Caffeine
750 UGML CAFFEINE.lcd 273nm,4nm
1000 µg/ml Caffeine
1 MGML CAFFEINE.lcd 273nm,4nm
mAU
min

Figure S2

A

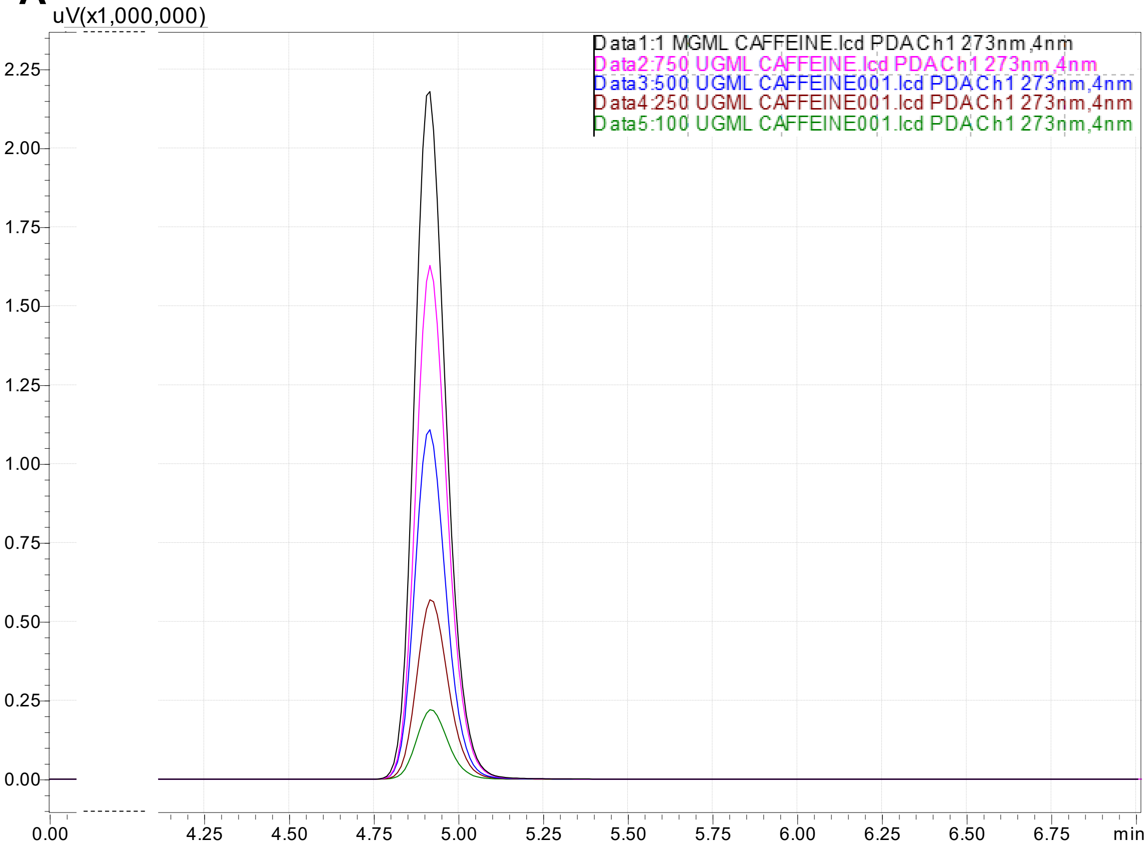


B

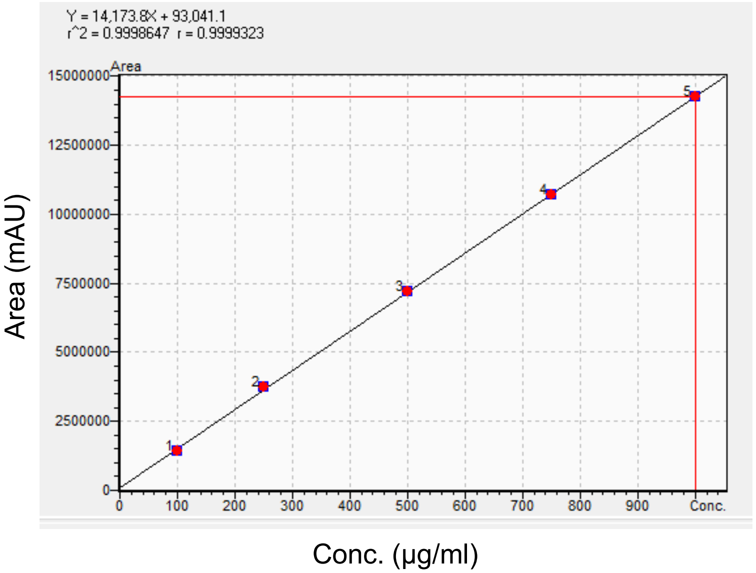


C

| Sample | Caffeine (µg/ml) | Area (mAU) |
|---|---|---|
| 1. | 100 | 1438296 |
| 2. | 250 | 3724623 |
| 3. | 500 | 7195397 |
| 4. | 750 | 10687825 |
| 5. | 1000 | 14271027 |

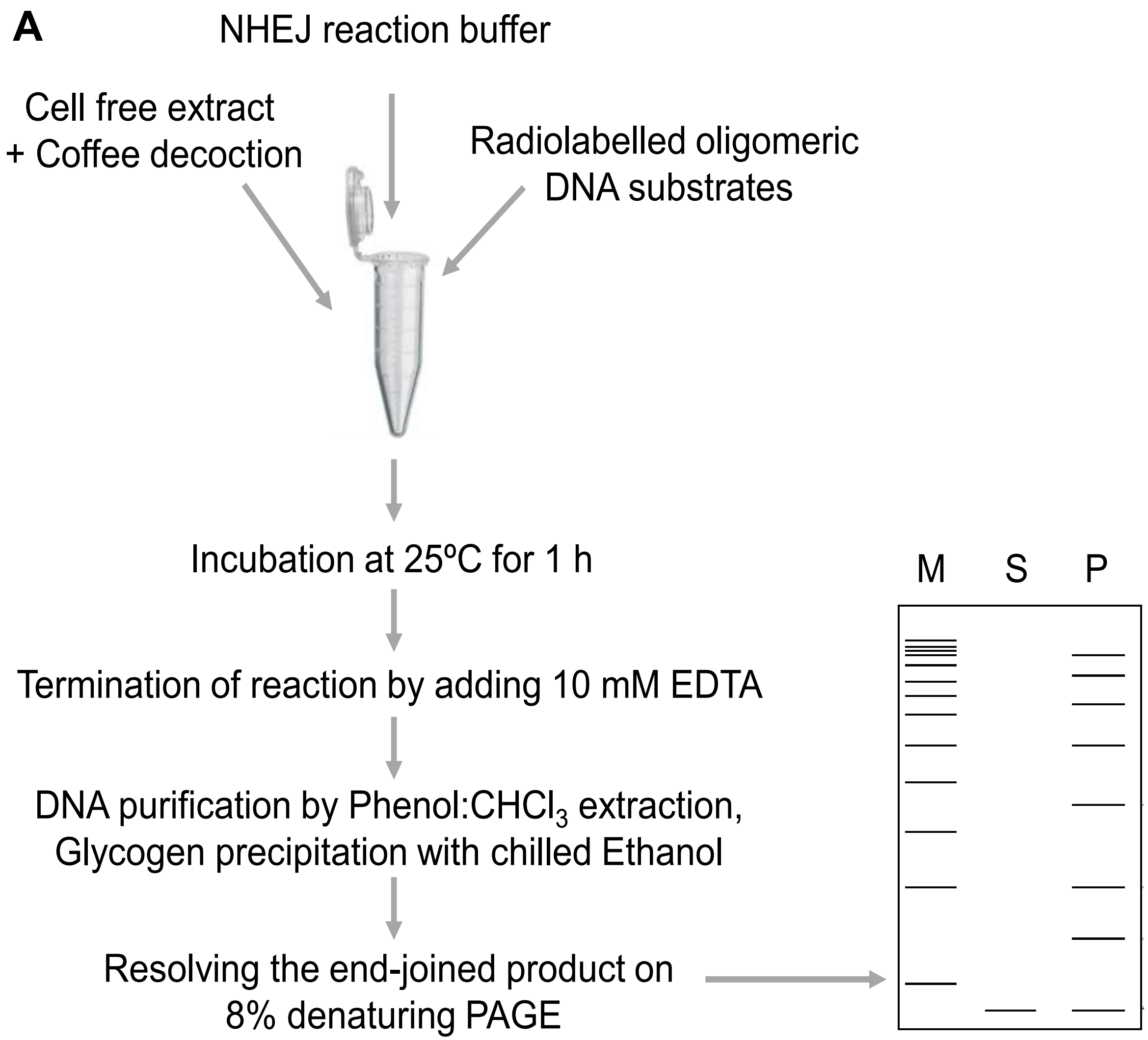
A
NHEJ reaction buffer
Cell free extract
+ Coffee decoction
Radiolabelled oligomeric
DNA substrates
Incubation at 25ºC for 1 h
Termination of reaction by adding 10 mM EDTA
DNA purification by Phenol:CHCl3 extraction,
Glycogen precipitation with chilled Ethanol
Resolving the end-joined product on
8% denaturing PAGE
M
S
P

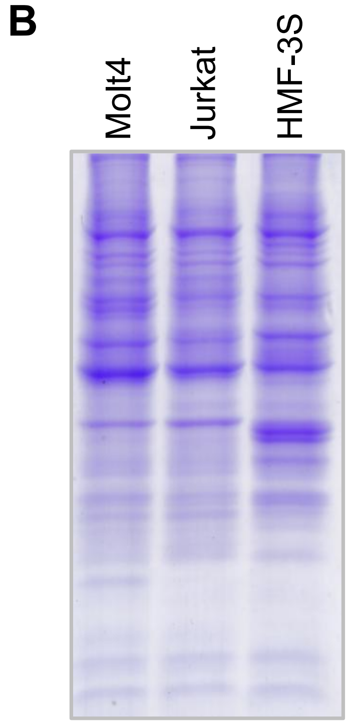
B
Molt4
Jurkat
HMF-3S

Figure S4

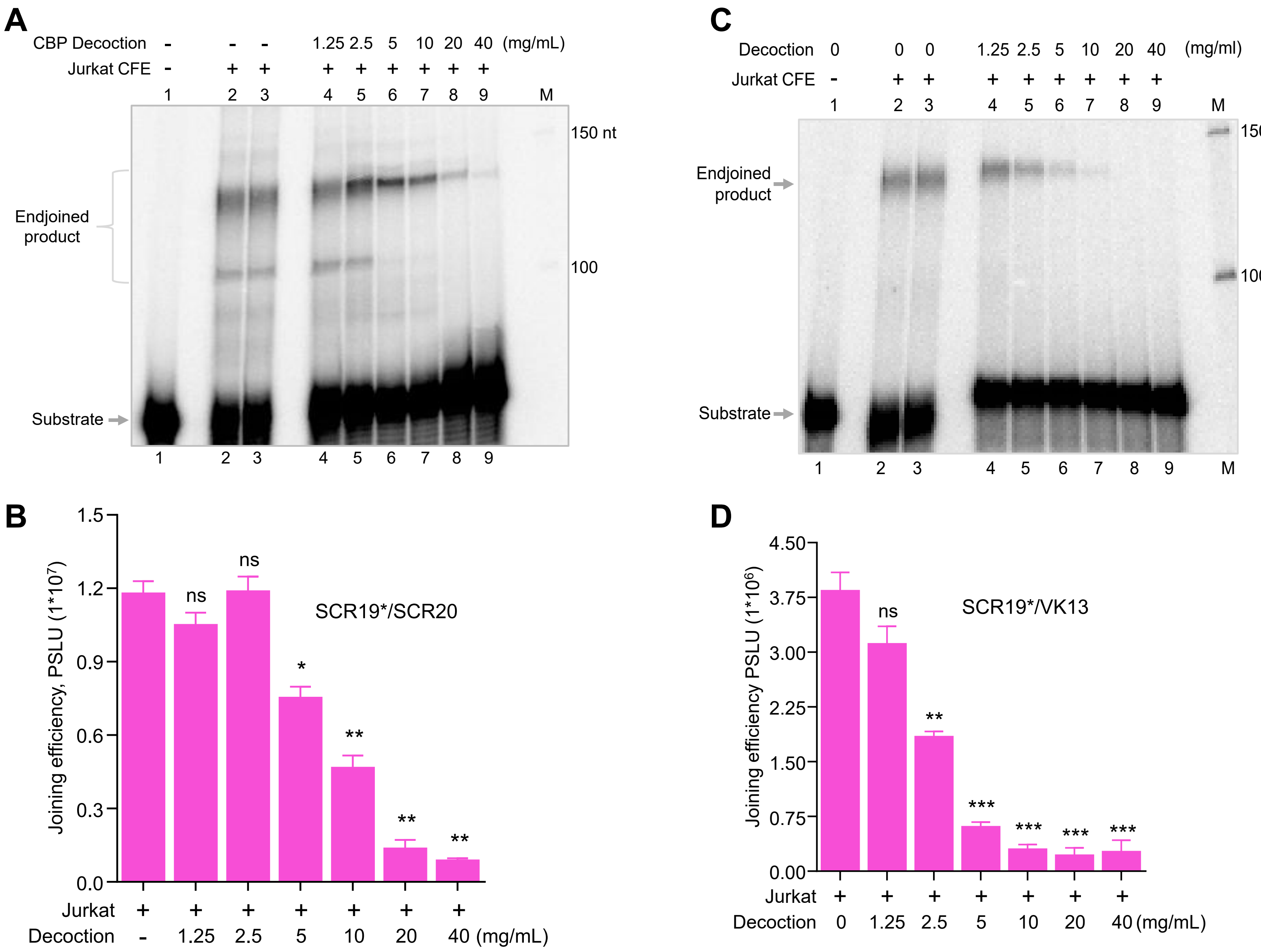

A
CBP Decoction - - - 1.25 2.5 5 10 20 40 (mg/mL)
Jurkat CFE - + + + + + + + +
1 2 3 4 5 6 7 8 9 M
150 nt
100
Endjoined product
Substrate
B
Joining efficiency, PSLU (1*10^7)
1.5
1.2
0.9
0.6
0.3
0.0
ns
ns
*
**
**
**
SCR19*/SCR20
Jurkat + + + + + + +
Decoction - 1.25 2.5 5 10 20 40 (mg/mL)
C
Decoction 0 0 0 1.25 2.5 5 10 20 40 (mg/ml)
Jurkat CFE - + + + + + + + +
1 2 3 4 5 6 7 8 9 M
150 nt
100
Endjoined product
Substrate
D
Joining efficiency PSLU (1*10^6)
4.50
3.75
3.00
2.25
1.50
0.75
0.00
ns
**
***
***
***
***
SCR19*/VK13
Jurkat + + + + + + +
Decoction 0 1.25 2.5 5 10 20 40 (mg/mL)

Figure S5

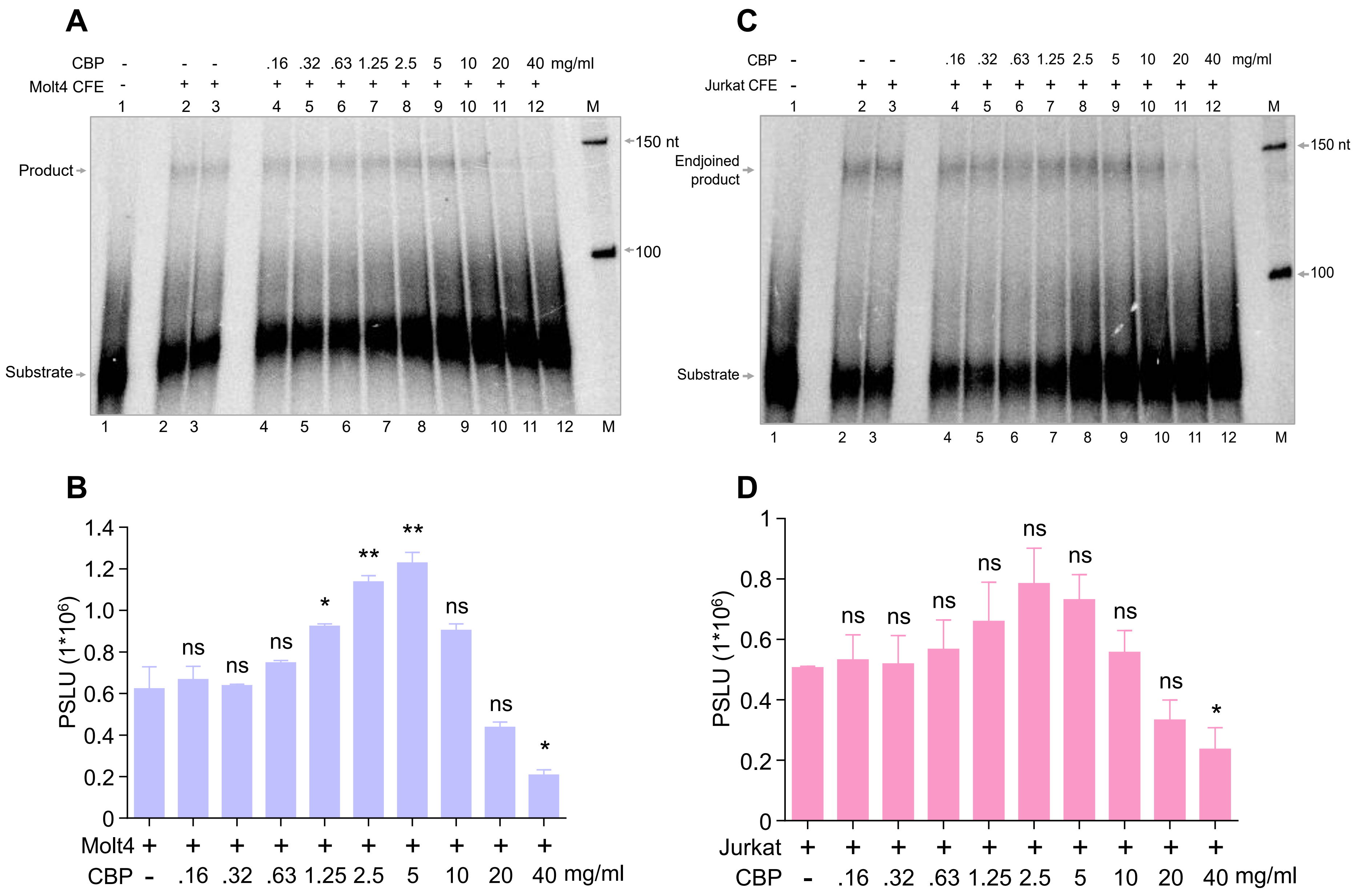

A
CBP - - - .16 .32 .63 1.25 2.5 5 10 20 40 mg/ml
Molt4 CFE - + + + + + + + + + + +
1 2 3 4 5 6 7 8 9 10 11 12 M
Product
Substrate
150 nt
100
C
CBP - - - .16 .32 .63 1.25 2.5 5 10 20 40 mg/ml
Jurkat CFE - + + + + + + + + + + +
1 2 3 4 5 6 7 8 9 10 11 12 M
Endjoined product
Substrate
150 nt
100
B
PSLU (1*10^6)
1.4
1.2
1.0
0.8
0.6
0.4
0.2
0
ns ns ns * ** ** ns ns *
Molt4 + + + + + + + + + +
CBP - .16 .32 .63 1.25 2.5 5 10 20 40 mg/ml
D
PSLU (1*10^6)
1
0.8
0.6
0.4
0.2
0
ns ns ns ns ns ns ns ns *
Jurkat + + + + + + + + + +
CBP - .16 .32 .63 1.25 2.5 5 10 20 40 mg/ml

Figure S6

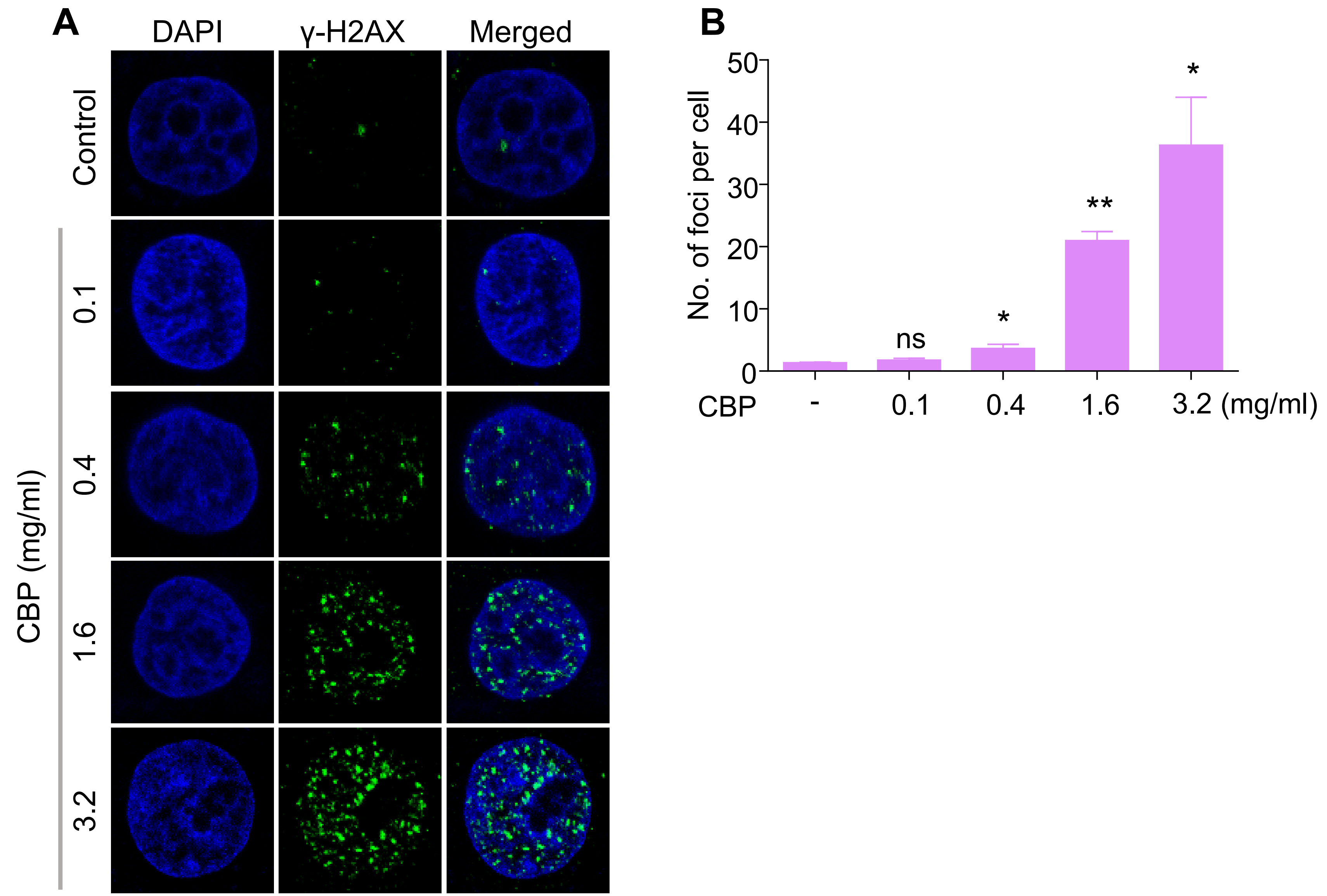
A
DAPI
γ-H2AX
Merged
Control
0.1
0.4
1.6
3.2
CBP (mg/ml)
B
No. of foci per cell
50
40
30
20
10
0
ns
*
**
*
CBP
-
0.1
0.4
1.6
3.2 (mg/ml)

Figure S7

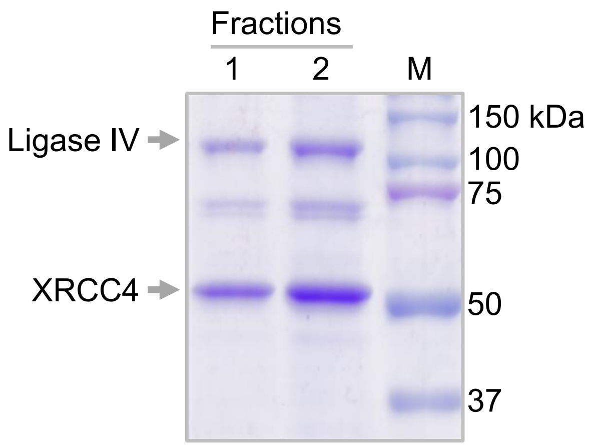
